# A spatial-temporal atlas of human islet pathophysiology identifies a size-dependent trajectory from compensation to decompensation

**DOI:** 10.64898/2026.01.29.701913

**Authors:** Tengli Liu, Rui Liang, Lanqiu Zhang, Wenmiao Ma, Xiangyu Wang, Huixia Ren, Shusen Wang

## Abstract

Type 2 diabetes (T2D) is characterized by progressive islet dysfunction, yet the transition from functional adaptation to failure remains poorly defined within the native tissue architecture. Using multiplex imaging mass cytometry, we systematically analyzed human pancreatic islets across a spectrum of non-diabetic (ND), prediabetic (PreD), and T2D donors, revealing that pathological remodeling is profoundly size-dependent. This remodeling is manifested as a coordinated evolution of subpopulation abundance, endocrine cell proportions, structural integrity, and protein expression profiles, revealing that islets of different sizes undergo divergent fates during disease progression. We identified a size-dependent vulnerability spectrum where medium-and large-sized islets (>100 μm in diameter) serve as the primary histopathological correlates of glycemic failure (HbA1c), exhibiting early density loss and structural disintegration. In contrast, small islets (30-100 µm in diameter) exhibited compensatory hormone upregulation during PreD. Notably, diverging from the ‘death of the small’ paradigm in Type 1 Diabetes, our data reveal a significant expansion of extra-islet endocrine clusters (EECs) that initiates during the compensatory stage, effectively preceding the onset of overt hyperglycemia. Finally, t-SNE clustering reconstructed a continuous trajectory of islet remodeling, capturing the phenotypic evolution of islets from normoglycemia through compensation to clinical decompensation. This study provides a high-resolution atlas of islet pathophysiology, offering new insights into T2D progression.

**Highlights:**

1. Islet remodeling during T2D progression is critically dependent on islet size.
2. Medium and large islets density negatively correlates with HbA1c.
3. Small islets exhibit resilience to metabolic stress during T2D progression.
4. EECs expansion suggests a process of α-cell-biased neogenesis.
5. t-SNE maps the islet trajectory from compensation to decompensation.

## INTRODUCTION

Type 2 diabetes (T2D) is a global epidemic characterized by the progressive failure of pancreatic islets to sustain sufficient insulin secretion to counteract peripheral insulin resistance ^1,2^, yet the cellular and structural basis underlying the transition from adaptive compensation to irreversible decompensation remains poorly defined. A fundamental and unresolved question persists: within the heterogeneous landscape of the human pancreas, why do some islets maintain functional resilience in the face of metabolic stress, while others succumb to structural disintegration and cellular dysfunction?

Human pancreatic islets display remarkable size heterogeneity, ranging from single endocrine cells and small clusters to large islets containing thousands of cells ^3,4^. This heterogeneity is not merely morphological; emerging evidence indicates that islet size directly dictates cellular composition, architecture, and consequent paracrine signaling dynamics ^3-10^. Yet in studying islets pathogenic alterations in diabetes progression, early studies preferred to focus on a single size class or treat islets as a homogeneous population in studying their pathogenic changing tread in diabetes progression, failing to systematically track size-dependent functional trajectories across the full spectrum of disease progression. Recent investigations have begun to address this issue and yielded breakthrough insights in type 1 diabetes (T1D)—uncovering previously unrecognized phenomena such as the high abundance of small islets and their early loss during disease initiation^11-13^. In contrast, analogous systematic analyses encompassing the full islet size spectrum and complete T2D progression trajectory remain lacking, leaving the size-dependent pathogenic mechanisms of T2D islet dysfunction uncharacterized. Moreover, beyond β-cell failure, accumulating evidence underscores the pathogenic role of non-β cells in T2D: hyperglucagonemia driven by α-cell dysfunction directly exacerbates hyperglycemia ^14-16^, while α-cell-derived glucagon and δ-cell-derived somatostatin also modulate both β-cell activity and glucose homeostasis ^17-22^. Yet, how the islet cellular composition dynamically shifts across different islet size classes during T2D progression remains unexplored.

To address these gaps, we leverage imaging mass cytometry (IMC), a powerful multiplexed imaging technology designed for in-depth analysis of complex tissues ^23-25^, to generate a high-resolution spatial-temporal atlas of over 5,300 human islets spanning non-diabetic, normoglycemic compensatory, prediabetic, and T2D states. Our 19-marker panel enables simultaneous profiling of endocrine cell identities, structural integrity, and functional markers at single-cell resolution, while strict matching of clinical covariates (age, BMI, lipid levels) ensures robust comparisons across groups. This study reveals a size-dependent trajectory of islet remodeling that redefines our understanding of T2D pathophysiology, uncovering both vulnerable islet populations that drive disease progression and resilient subsets with therapeutic potential.

## RESULTS

### Study overview

While previous research has documented the aggregate loss of β-cell mass, the precise spatial dynamics and structural evolution across different islet size cohorts—particularly the nascent extra-islet endocrine clusters (EECs)—remain poorly defined.

To investigate the structural and molecular evolution of human islets across the full-size continuum, we utilized IMC to analyze a pancreas cohort from the Human Islet Resource Center (China), including 10 ND, 8 PreD, and 12 T2D cases matched for age, sex, and BMI (Figure 1A&1B). Because regional heterogeneity can significantly skew islet composition data, we exclusively analyzed sections from the pancreatic tail, selecting 245 regions of interest (ROIs) that encompassed 5,313 islets and approximately 1.7 million single cells within islets (Figure 1C).

**Figure 1.**
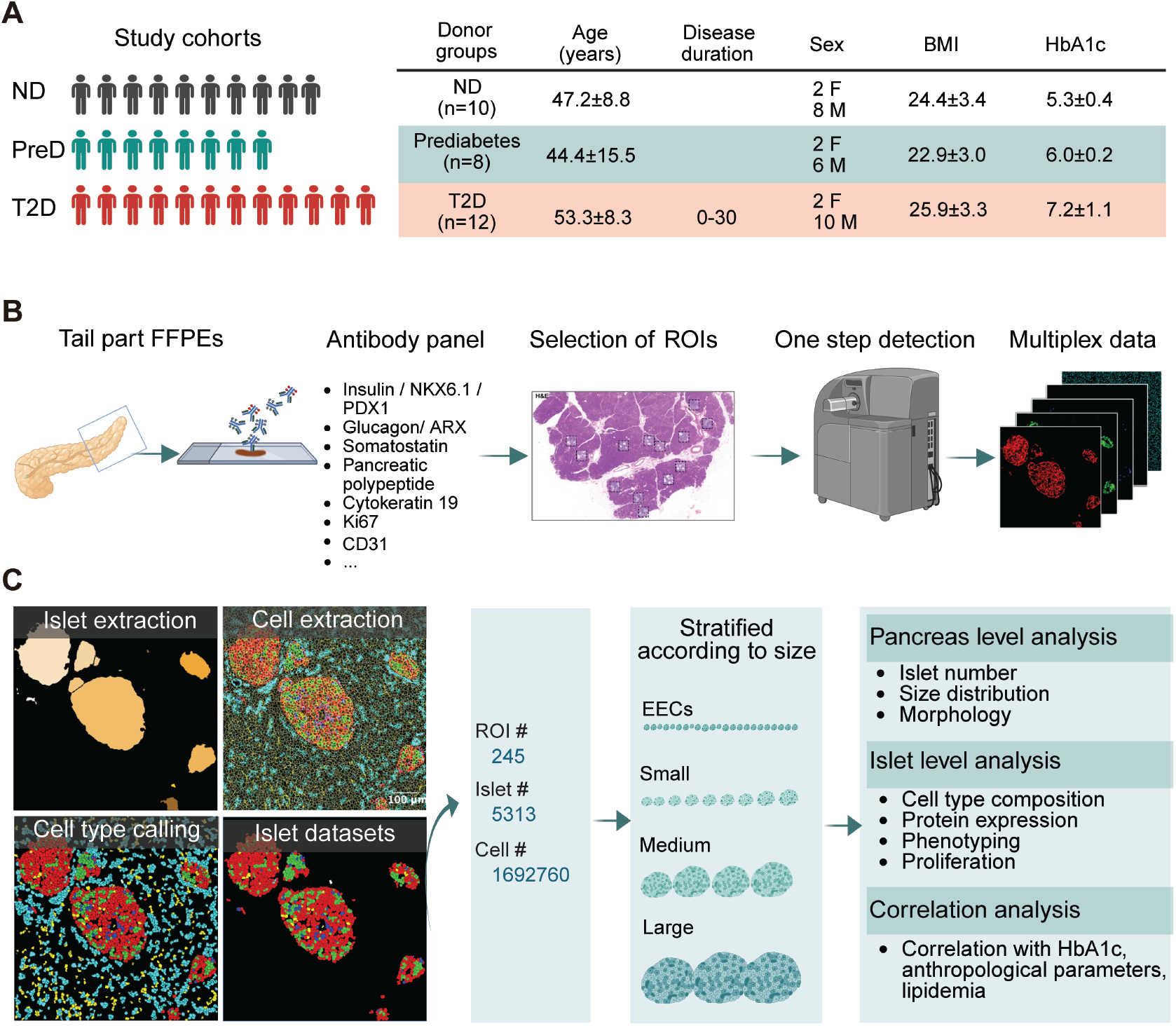
Workflow for imaging mass cytometry study of human pancreases. (A) Donor characteristics. (B) Schematic overview of the experimental platform. Paraffin sections from the tail region of human pancreas samples obtained from organ donors were stained with metal isotope-labeled antibodies. A laser sequentially ablates tissue spots, and the ablated material is analyzed by mass cytometry to identify distinct metal isotope labels. Based on the metal isotope composition at each spot, an image of the ablated area is reconstructed for each measured marker. (C) Data analysis pipeline. Cellular segmentation and protein expression analysis were performed using imaging mass cytometry (IMC) data. Subsequently, islet segmentation and cell-type annotation were conducted based on the derived cellular and protein masks, resulting in an islet-specific single-cell dataset. Green, insulin; Red, glucagon; Blue, somatostatin (SST); Gray, pancreatic polypeptide (PP); Cyan, cytokeratin 19 (CK19); Yellow, CD34. Scale bar: 100 µm.

We employed an integrated 19-protein IMC panel that allowed us to simultaneously co-register markers for all five endocrine cell types, key transcription factors (PDX1, NKX6.1, ARX), pancreatic ductal cells (cytokeratin 19, PDX1), endothelial and stromal cells (CD34, α-SMA), cell proliferation markers (Ki67, phosphorylated histone H3), as well as cytoskeletal and cell membrane markers (β-actin and CD99) (Figure 1B).

Following rigorous antibody validation, we implemented a robust analytical pipeline, utilizing CellProfiler for single-cell segmentation and hormone-based masks for islet delineation to extract multiplex spatial protein expression data from each individual cell (Figure 1C). These islets were further stratified according to their size and changing trends during T2D progression, which enables us to move beyond aggregate mass measurements to a high-resolution view of the islet architectural remodeling during T2D progression (Figure 1C).

### Size-dependent pathogenic remodeling of the human islet landscape during diabetes progression

To characterize islet morphological remodeling across diabetes progression, we partitioned all islets into 26 exponentially spaced bins based on their area—where the width of each area interval expands exponentially with increasing numerical value. The projection of the area and corresponding diameter for each bin is presented in Figure S4. Then based on their enrichment patterns and dynamic alterations during T2D progression, the 26 bins were subsequently subdivided into four size classes: extra-islet endocrine clusters (EECs, with mean diameters <30 μm), small (30–100 μm), medium (100–185 μm), and large (>185 μm) islets (Fig. 2A). Representative IMC staining images (INS/GCG/SST/DNA merge) illustrated the size heterogeneity of islets across groups.

**Figure 2.**
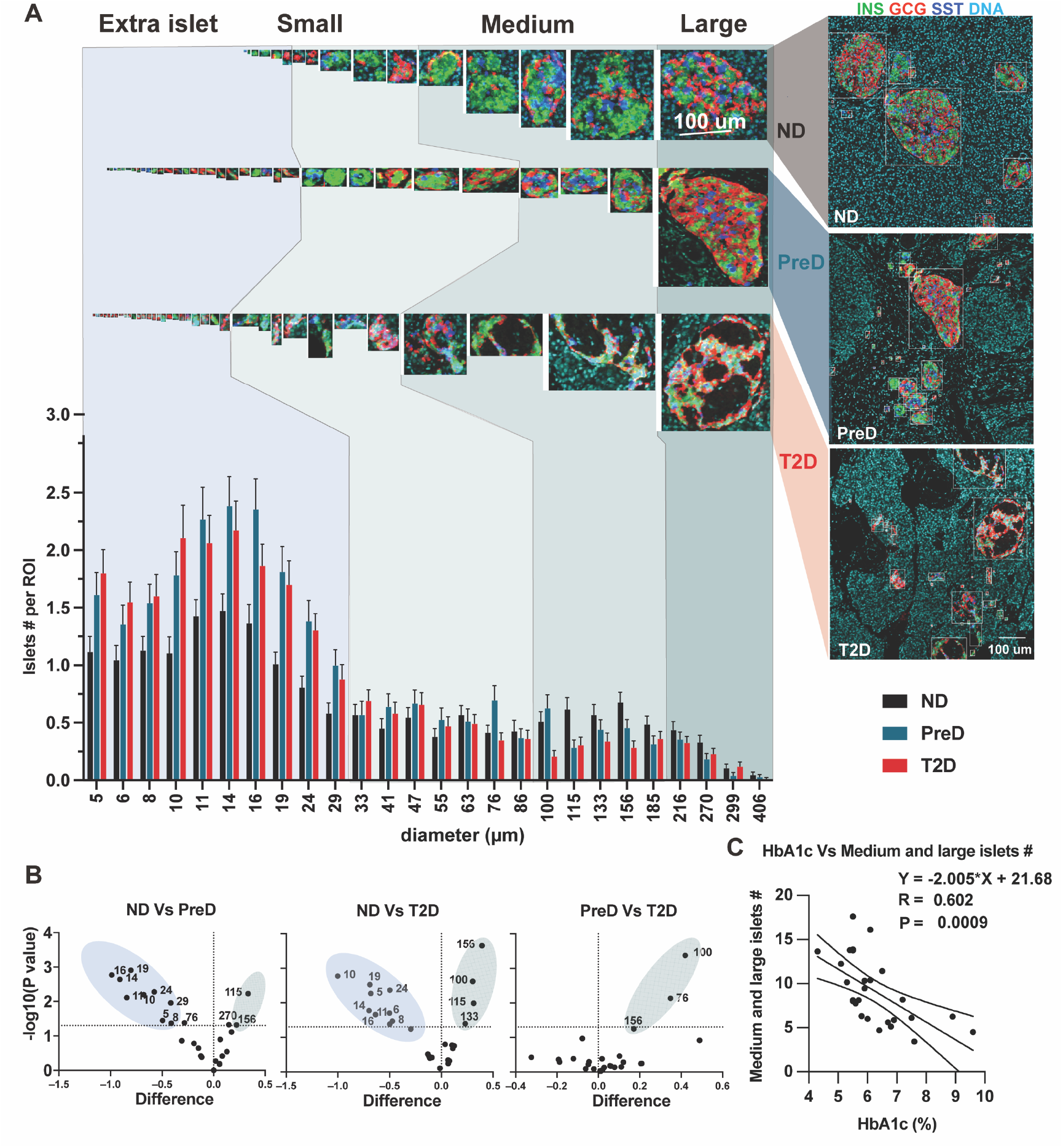
Remodeling of human islet size distribution across nondiabetic, prediabetic, and type 2 diabetic stages. (A) Islet count per region of interest (ROI), stratified by exponentially binned area ranges. Bin labels indicate the mean diameter for each bin. Islets were classified into four size categories: extra-islet endocrine clusters (EECs, with mean diameter < 30 μm), small (with mean diameter of 30-100 μm), medium (with mean diameter > 100 μm), and large (with mean diameter >200 μm). Representative IMC images islets (co-stained for insulin (INS), glucagon (GCG), somatostatin (SST), and DNA) of each size class from ND, PreD, and T2D ROIs, are displayed above the column chart. (B) Volcano plots of statistical comparisons between group pairs (ND vs PreD, ND vs T2D, PreD vs T2D): x-axis = log2-fold change in islet count per bin; y-axis = -log10(P-value). Dots in the upper-right/left quadrants denote bins with statistically significant changes in islet abundance. (C) Correlation analysis between HbA1c levels and the number of medium and large islets across all subjects.

Multiplexed IMC revealed a striking redistribution of the endocrine landscape as disease progressed. Quantitative analysis per ROI identified a progressive shift toward smaller islet dimensions: EECs significantly accumulated in the PreD and T2D pancreas, whereas medium and large islets exhibited a steady decline in frequency (Figure 2A). This observation reveals a notable “extra-islet paradox” in T2D; unlike the “death of the small” paradigm seen in type 1 diabetes, the T2D environment is characterized by an active expansion of small endocrine units (Figure 2A&3A), likely representing a compensatory neogenic response.

**Figure 3.**
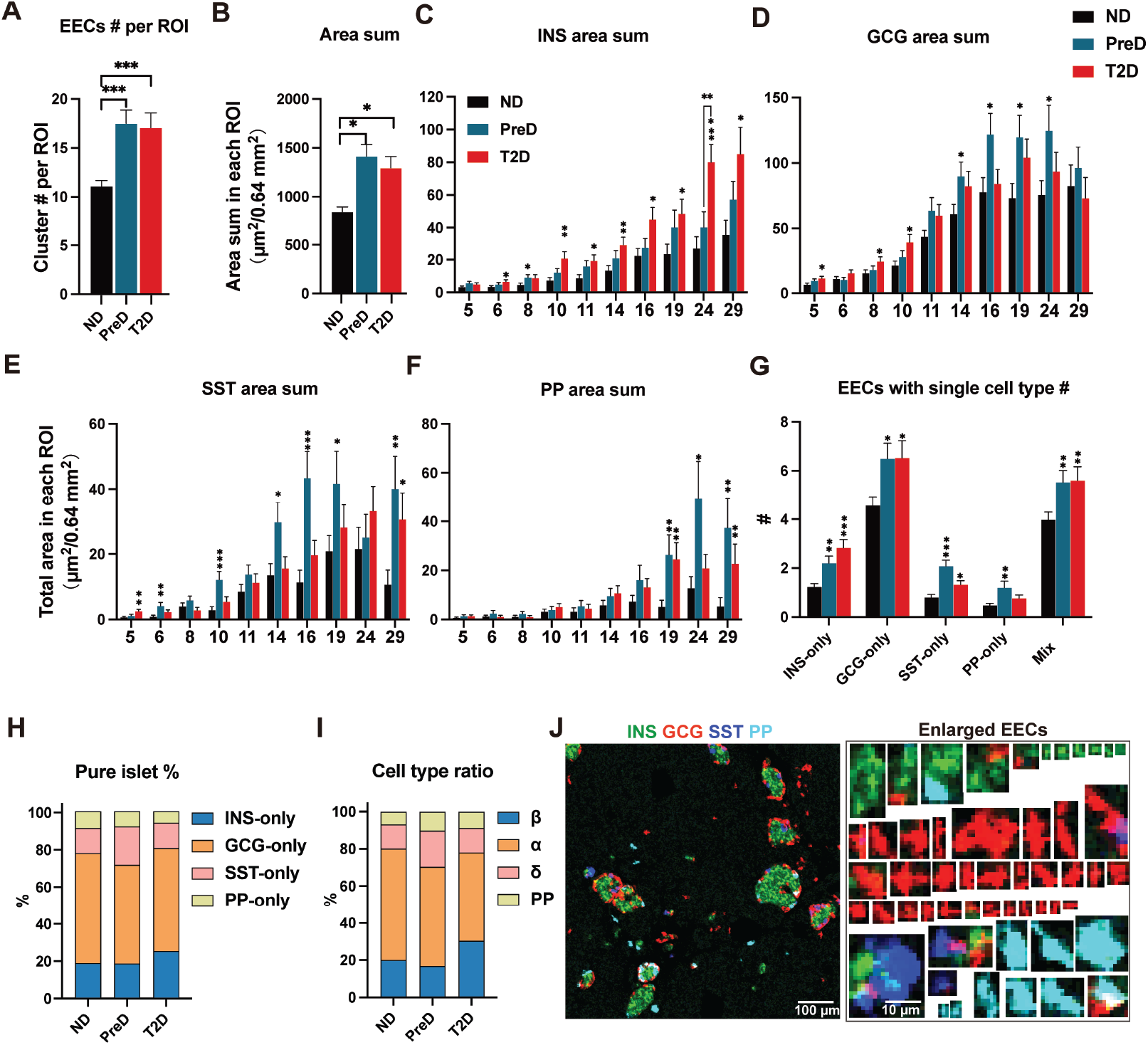
Expansion of α cell-dominant extra-islet endocrine clusters across nondiabetic, prediabetic, and type 2 diabetic pancreases. (A) Counts of extra-islet endocrine clusters (EECs) per ROI. (B) The cumulative area of EECs per ROI in each group. (C-F) The cumulative insulin-positive area (C), glucagon-positive area (D), somatostatin-positive area (E), and pancreatic polypeptide (PP)-positive area (F) of EECs per ROI in each group. (G) Counts of EECs expressing only one hormone: INS-only, GCG-only, SST-only, or PP-only. (H) Proportion of EECs composed of a single hormone type (I) Proportion of each cell type (α, β, δ, and PP cells) in all cells within EECS per ROI. (J) Representative IMC images of EECs (co-stained for INS, GCG, SST, and PP), Scale bar = 100 μm; enlarged insets highlight the morphology of hormone-specific EECs, Scale bar = 10 μm. Data are mean ± SEM; ^*^P < 0.05, ^**^P < 0.01, ^***^P < 0.001.

To statistically define the “tipping point” of metabolic failure, we utilized volcano plot to display the statistical difference in islet size distributions across clinical stages. Comparisons between ND and PreD donors identified significant increase in EECs and decreases in medium (115 and 156 µm in diameter) and large-sized (270 in diameter) islets, confirming that structural remodeling initiates during the subclinical stage (Figure 2B, left). Correlation analysis revealed a robust negative correlation between medium and large islet abundance and HbA1c levels (Figure 2C). These reulsts suggest that the maintenance of the medium and large islets population (>100 μm) is the primary histopathological determinant of systemic glycemic stability.

### Expansion and α-cell predominance of extra-islet endocrine clusters during T2D progression

Consistent with increased EEC abundance, the relative contribution of EECs to total islet area was also heightened in PreD and T2D compared to ND (Figure 3B), reflecting the growing morphological prominence of EECs as the endocrine landscape is reshaped.

A detailed mapping of endocrine cell composition within these clusters revealed divergent, size-specific remodeling patterns. In PreD donors, the significant increase was primarily restricted to single cells, yet in T2D donors, the insulin-positive (INS+) area per ROIwas significantly elevated across nearly all EEC size ranges, suggesting significant expansion occurred until T2D (Figure 3C). Conversely, the glucagon-positive (GCG+) area followed a different trajectory: in PreD, the increase was predominantly localized to larger EEC subsets (14–24 μm), yet in T2D, a significant expansion occurred only in the smallest clusters (≤10 μm). Similarly, somatostatin-positive (SST+) area was significantly increased across most EEC sizes in the PreD stage, but not in T2D. This may suggest an early, widespread compensatory response that becomes more restricted as the disease progresses to overt T2D.

A defining characteristic of the extra-islet landscape was the high prevalence of “pure” endocrine clusters; over 65% of identified EECs consisted of cells expressing only a single hormone type (Figure S5). Among these single-hormone units, pure GCG+ (α-cell only) islets were the most abundant subtype (Figure 3G and 3H). Quantitative cell-type ratio analysis further confirmed this pattern, identifying α-cells as the predominant cell type within the extra-islet compartment (Figure 3I). Representative IMC images (INS/GCG/SST/PP) visualized these EECs as discrete, hormone-specific clusters characteristic of the diseased pancreas (Figure 3J). Collectively, these data demonstrate that the T2D environment triggers a robust expansion of EECs, likely through compensatory neogenesis, with a distinct developmental bias toward the α-cell lineage.

### Islet endocrine composition and hormone expression profiles

To investigate how islet size influences the pathological transformation of the endocrine pancreas, we performed a stratified analysis of small, medium, and large islets (Figure 4A). EECs were not included here since most extra islets consisted of a single endocrine cell type. The cumulative area of each sized islets per ROI revealed a significant and progressive reduction in the total area of medium and large islets in both PreD and T2D donors, while that of small islets remained relatively stable across all groups (Figure 4B). Consistently, INS+, GCG+, or SST+ area sum per ROI exhibited a congruent decreasing trend, whereas PP+ area sum showed no significant change (Figure 4B).

**Figure 4.**
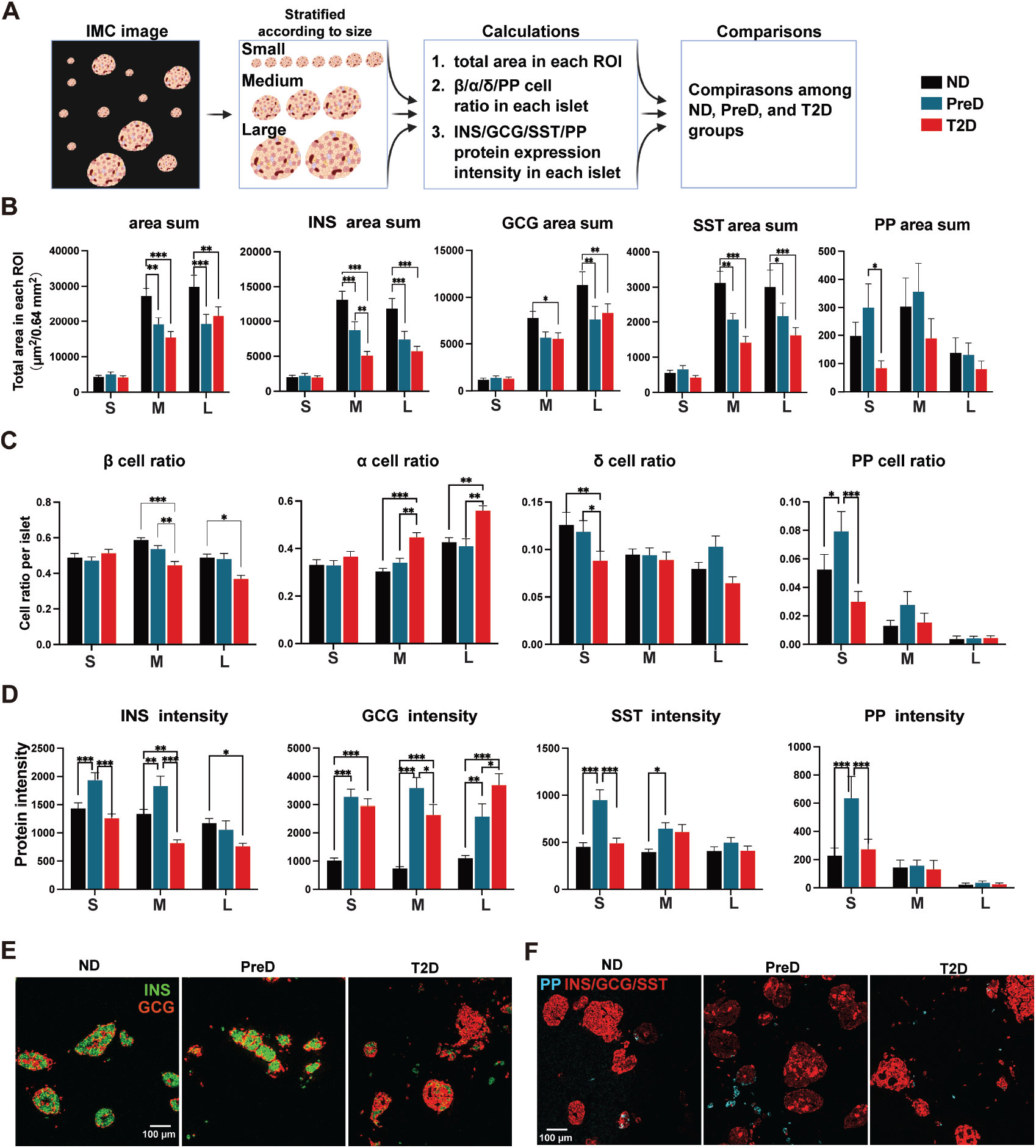
Size-stratified analysis of islet area, cellular composition, and hormone protein expression in human pancreatic islets across ND, PreD, and T2D groups. (A) Schematic workflow of the analytical pipeline: Islets were stratification into Small (S), Medium (M), and Large (L) size classes. Three key metrics were calculated: 1) total islet area per ROI; 2) β/α/δ/ PP cell ratios per islet; 3) protein intensity of insulin (INS), glucagon (GCG), somatostatin (SST), and PP per islet. These metrics were subsequently compared across ND, PreD, and T2D groups. (B) Bar plots depicting (from left to right) total islet area per ROI, and summed area of INS, GCG, SST, and PP in S, M, L islets across ND, PreD, and T2D groups. (C) Cellular composition profiles of size-stratified islets: β cell ratio, α cell ratio, δ cell ratio, and PP cell ratio (from left to right) in S, M, L islets across ND, PreD, and T2D groups. (D) Protein intensity of endocrine hormones (INS, GCG, SST, PP; from left to right) in S, M, L islets across ND, PreD, and T2D groups. (E) Representative IMC images showing INS (red) and GCG (green) in islets from ND, PreD, and T2D groups; scale bar = 100 μm. (F) Representative IMC images of PP (cyan)-positive islets in ND, PreD, and T2D groups; Red: INS/GCG/SST; scale bar = 100 μm. Data are mean ± SEM; ^*^P < 0.05, ^**^P < 0.01, ^***^P < 0.001.

A detailed assessment of cellular composition showed that the proportion of β-cells significantly decreased in medium and large islets during the transition to T2D, accompanied by a notable increase in the α-cell ratio (Figure 4C and 4E). Interestingly, these cellular ratios remained largely unchanged during the PreD stage for larger islets (Figure 4C and 4E). In contrast, small islets exhibited alterations in non-β endocrine cells, characterized by a pronounced increase in the PP-cell ratio in the PreD stage (Figure 4C and 4F), followed by a significant reduction in the δ-cell ratio in T2D (Figure 4C).

We further quantified hormone expression intensity to resolve the functional states of these size-specific cohorts (Figure 4D). PreD islets exhibited a robust compensatory signature, with significantly elevated levels of INS, GCG, SST, and PP, a phenomenon most prominent in small islets. However, as disease progressed to T2D, islets entered a decompensated state, marked by a decline in INS expression within medium and large islets, while GCG expression increased across all size classes (Figure 4D). Collectively, these data demonstrate the temporal size-dependent remodeling of different endocrine cell types in density, cell proportion and hormone content in islets, highlighting the transient insulin compensation in PreD and overall α-cell hyperplasia in PreD and T2D.

### Islet architectural integrity deteriorates across diabetes progression

To assess structural disruption of islets during diabetes development, we quantified key morphological parameters (reflecting shape irregularity and structural compactness) in small, medium, and large islets from ND, PreD, and T2D pancreases (Figure 5A–5D).

**Figure 5.**
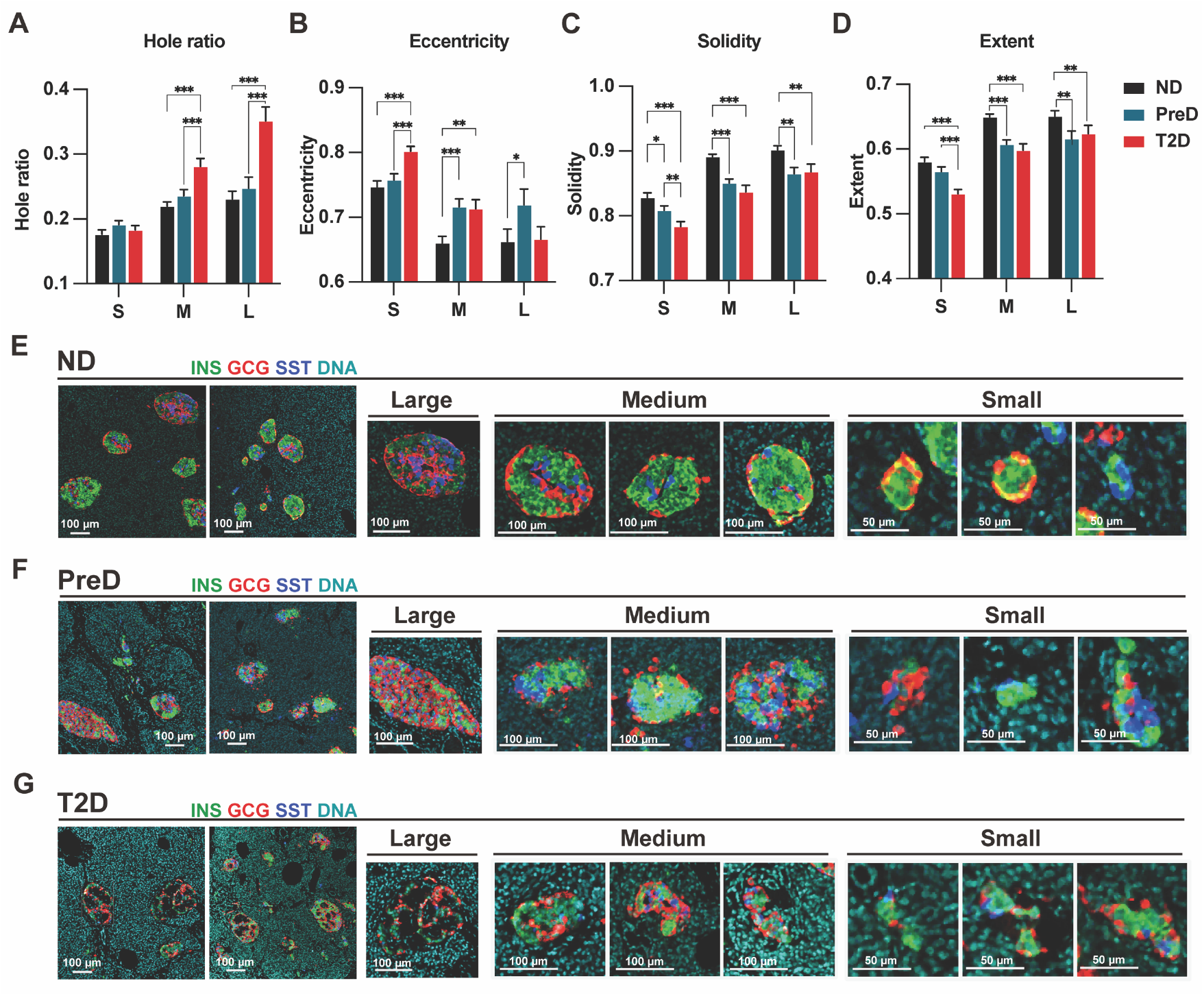
Progressive architectural disruption of human islets across ND, PreD, and T2D stages. (A–D) Quantitative analysis of islet morphological parameters (stratified by small (S), medium (M), and large (L) islet size classes) in ND, PreD, and T2D pancreases: Hole ratio (measure of intra-islet void density, A), Eccentricity (deviation of islet shape from a circular profile, B), Solidity (index of islet structural compactness, C), Extent (spatial spread of islet tissue, D). Data are mean ± SEM; ^*^P < 0.05, ^**^P < 0.01, ^***^P < 0.001. (E–G) Representative IMC images of islets (co-stained for INS, GCG, SST, and DNA) across groups: ND islets (exhibiting compact, regular morphology, E), PreD islets (mild structural irregularity, F), T2D islets (prominent intra-islet voids, fragmented contours, and distorted boundaries, G). Scale bars = 100 μm (main panels) and 50 μm (small-islet insets).

For hole ratio (a measure of intra-islet voids), medium and large islets exhibited the most striking changes: medium and large islets in T2D showed a significant increase in hole ratio relative to their ND and PreD counterparts, indicating enhanced intra-islet cavitation in advanced disease (Figure 5A) Furthermore, Hematoxylin and Eosin (H&E) staining suggested these voids likely represented amyloid deposits (Figure S6).

Islet eccentricity (a metric of deviation from a circular profile) was significantly increased in medium and large islets as early as the PreD stage. In contrast, small islets maintained their circularity in PreD and only exhibited a significant increase in eccentricity in the T2D stage, suggesting a delayed onset of shape distortion in this size class (Figure 5B).

Solidity (a marker of structural compactness) and extent (reflecting the spatial spread of islet tissue) both declined significantly in medium and large islets during the PreD and T2D stages, indicating an early and progressive loss of compact, organized architecture (Figure 5C & 5D). Notably, small islets preserved their extent in the PreD stage, with significant decreases observed only in the T2D stage. This pattern mirrors the delayed change in eccentricity, further supporting the relative structural resilience of small islets during the early phase of diabetes progression.

Representative imaging mass cytometry (IMC) images (INS/GCG/SST/DNA co-staining; Figure 5E–5G) visually confirmed these structural alterations: ND islets maintained compact, regular morphologies across all size classes (Figure 5E); PreD islets displayed mild irregularities, predominantly in the medium and large subsets (Figure 5F); and T2D islets exhibited widespread structural disruption, including prominent voids, fragmented boundaries, and distorted contours in large islets, alongside shape irregularities in small and medium islets (Figure 5G), consistent with the size-dependent and progressive nature of islet architectural deterioration observed in the quantitative analyses.

### tSNE clustering of medium/large islets identifies early islet remodeling driven by increasing lipids prior to glycemic elevation

To mitigate limitations inherent to cross-sectional human pancreatic tissue data, we sought to define trajectories of islet remodeling across diabetes progression by comprehensively profiling all medium and large islets from ND, PreD, and T2D groups. We applied t-distributed Stochastic Neighbor Embedding (tSNE) clustering to these islets, using area-normalized hormone expression levels as features; this analysis resolved 5 distinct, phenotypically discrete islet populations (color-coded in the tSNE map, Figure 6A & B). Representative IMC images (INS/GCG/SST/PP co-staining) visually validated the unique hormone expression profiles of each population (Figure 6C), confirming the distinctness of the tSNE-derived clusters.

**Figure 6.**
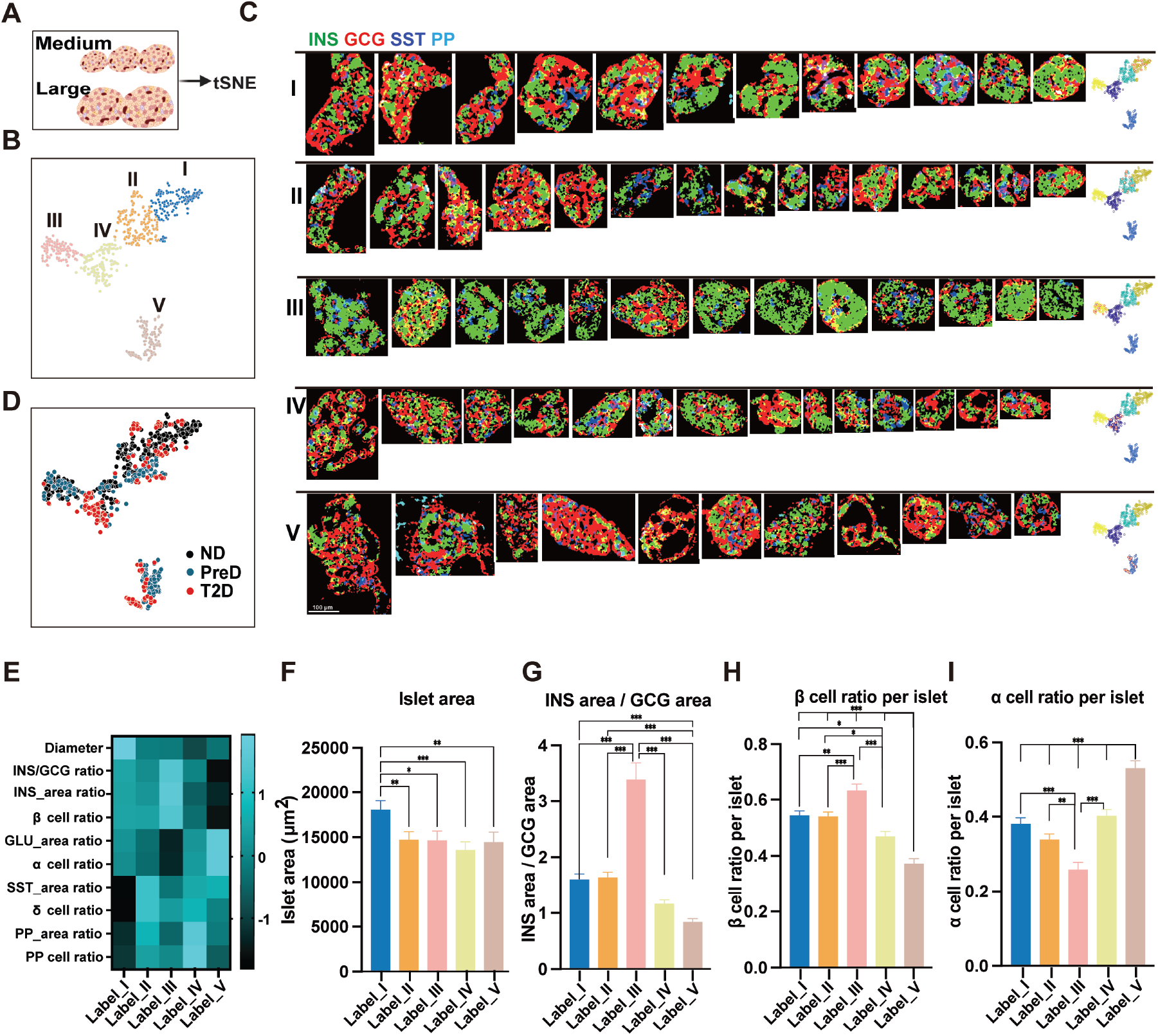
tSNE-based clustering of medium/large human islets identifies phenotypically distinct subpopulations. (A&B) Medium and large sized islets underwent t-distributed stochastic neighbor embedding (tSNE) dimentional reduction clustering based on islet area-normalized hormone expression levels, revealing five phenotypically distinct subpopulations (denoted as Label I–V)) (each dot represents one islet). (C) Representative IMC images of each subpopulation, displaying the expression patterns of INS (red), GCG (green), SST (blue), and PP (cyan). Scale bar = 100 μm. (D) tSNE plot colored by disease status (ND: black; PreD: blue; T2D: red), illustrating the enrichment profile of each subpopulation across ND, PreD, and T2D groups. (E) Heatmap summarizing key phenotypic features (islet diameter, INS/GCG ratio, cell area ratios, and cell number ratios for β/α/δ/PP cells) across populations labeled I–V. (F–I) Quantitative analyses across populations: Islet area (F); INS area/GCG area ratio (G); β cell ratio per islet (H); α cell ratio per islet (I). Data are presented as mean ± SEM; ^*^P < 0.05, ^**^P < 0.01, ^***^P < 0.001.

Profiling molecular and cellular features across these populations revealed a directional trajectory of islet phenotype shift. Population I exhibited larger islet size (Figure 6E and 6F). Insulin expression and β-cell proportions increased progressively from Populations I and II to Population III (reaching the highest levels in Population III), then declined sharply in Population IV, and dropped to the lowest in Population V (Figure 6E, G, H). Conversely, glucagon (GCG) expression and α-cell proportions followed the inverse pattern: decreasing from Populations I and II to III, then rising in Population IV, and peaking in Population V (Figure 6E, G, I). This opposing trend yielded Population III as the cluster with the highest INS-to-GCG area ratio, while Population V exhibited the lowest β-cell proportion and highest α-cell percentage (Figure 6G–I).

Islet distribution across clusters further aligned with disease stage (Figure 6D). ND islets were concentrated in Populations I, II, III (characterized by high INS expression area and β-cell ratio), with a few islets distributed in Population IV. PreD and T2D islets were enriched in Populations IV and V (marked by reduced β-cell ratio and elevated α-cell ratio). Notably, Population V— displaying the most severe β-cell loss and α-cell expansion—was exclusive to PreD and T2D donors; in contrast, Population III (the most β-cell-rich cluster) was primarily composed of ND and PreD islets (Figure 6D & 7A).

We next examined ND islet heterogeneity at the donor level: a heatmap of population distribution across individual ND samples revealed two distinct subgroups (ND_Gr1, ND_Gr2; Figure 7B). Representative IMC images of ROIs from these subgroups visually confirmed their distinct population profiles: ND_Gr1 ROIs were enriched in population I/II (classic ND islets), while ND_Gr2 ROIs contained more population III (compensatory islets; Figure 7C).

**Figure 7.**
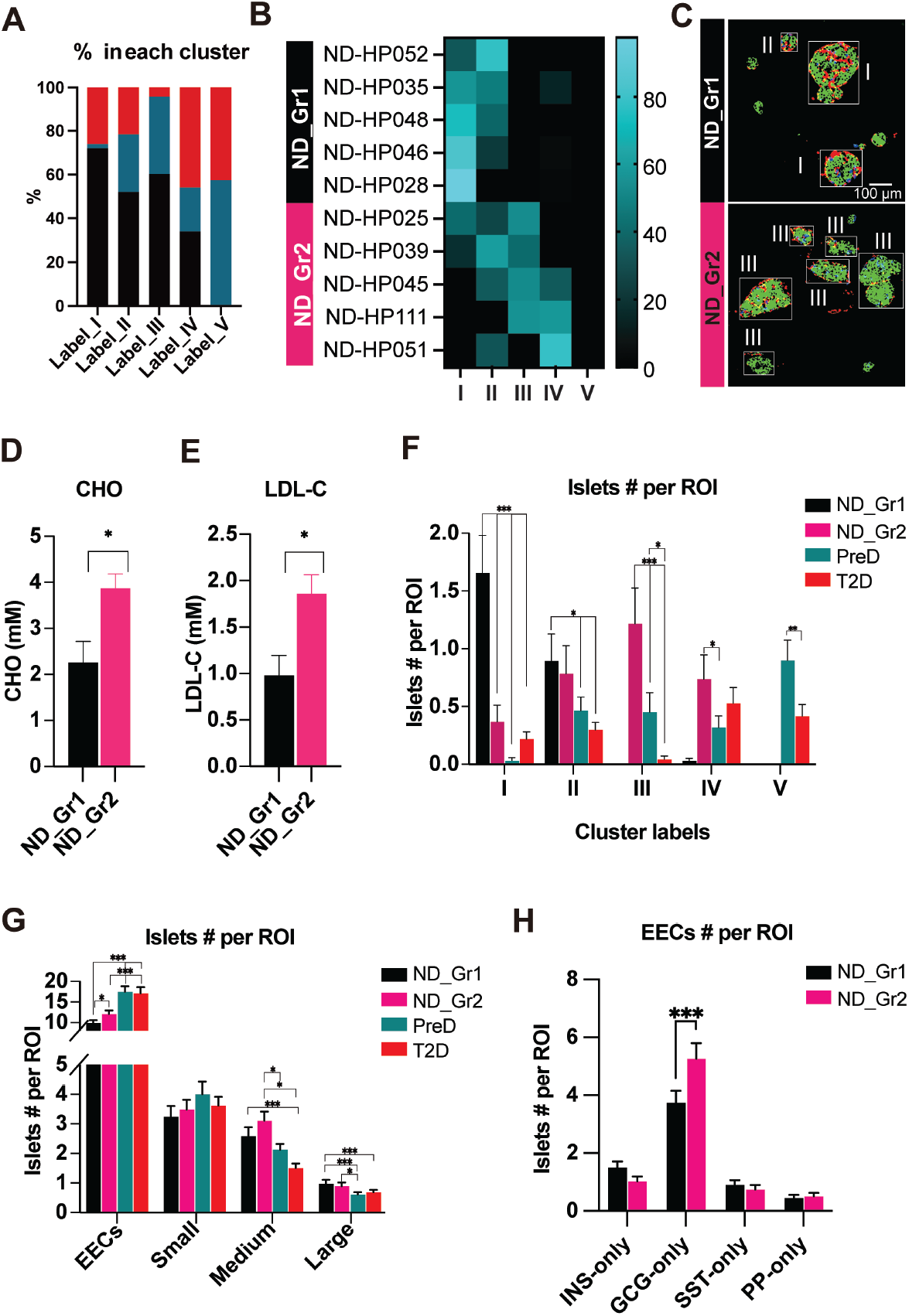
Distribution of tSNE-derived islet populations in ND, preD and T2D. (A) Percentage of islets from each disease stage (ND, PreD, T2D) within each of the five populations. (B) Heatmap showing the enrichment profile of islet populations across all ND samples, which are stratified into two subgroups (ND_Gr1 and ND_Gr2) based on subpopulation distribution patterns. (C) Representative IMC images of ROIs from ND_Gr1 and ND_Gr2; insets highlight islets corresponding to distinct subpopulations (Label I–III). Scale bar = 100 μm. (D&E) Cholesteron (CHO, mmol/L) levels and low-density lipoprotein cholesterol (LDL-C, mmol/L) levels of subjects in ND_Gr1 and ND_Gr2. (F) Number of islets per ROI for each population (Label I–V) across ND_Gr1, ND_Gr2, PreD, and T2D groups. (G) Number of islets per ROI across size classes (EECs, Small, Medium, Large) in ND_Gr1, ND_Gr2, PreD, and T2D groups. (H) Bar plot depicting the number of single-hormone EECs (INS-only, GCG-only, SST-only, PP-only) per ROI in ND_Gr1 and ND_Gr2. Data are mean ± SEM; ^*^P < 0.05, ^**^P < 0.01, ^***^P < 0.001.

Clinical parameter comparison between ND_Gr1 and ND_Gr2 showed no differences in HbA1c, age, triglycerides, or BMI, but ND_Gr2 exhibited significantly higher total cholesterol (CHO) and low-density lipoprotein cholesterol (LDL-C) levels (Figures 7D&7E). This suggests elevated lipids metabolic stress may drive early islet remodeling in normoglycemic individuals, shifting islets from the classic ND phenotype (Population I) to a compensatory state (Population III). This compensatory islet subtype sustained in PreD but lost in T2D, where enriched with α cell dominant population IV and V (Figure 7F).

Analysis of size-classed islets revealed that ND_Gr2 showed expanded, α-cell dominant EECs (Figure 7G and 7H)—though this expansion was less prominent than in the PreD and T2D stages— and also exhibited elevated medium islet counts (Figure 7G). These findings indicate that subjects in ND_Gr2 occupied an insulin secretory compensatory stage prior to prediabetes or any detectable glycemic elevation.

## DISCUSSION

This comprehensive spatial analysis of over 5,300 human islets using IMC provides a high-resolution atlas of pancreatic remodeling, revealing that the transition from metabolic health to T2D is defined by a size-dependent structural and cellular collapse, characterized by the attrition of the mdium and large sized islets and the expansion of micro-endocrine units. Our findings identify an islet evolution trajectory from physiological compensation to decompensation, providing a structural basis for the subclinical progression of glycemic failure.

### Pathogenic EEC expansion and α-cell-biased neogenesis

A fundamental divergence between type 1 and type 2 diabetes pathology lies in the fate of the smallest endocrine units, which in T2D undergo a robust, paradoxical expansion rather than autoimmune attrition. Recent high-resolution mapping of the type 1 diabetes (T1D) landscape has demonstrated that small endocrine objects, including extra-islet endocrine clusters (EECs), are highly susceptible to autoimmune destruction and are among the first units lost during disease progression^11^. In stark contrast, our data reveal that within the T2D milieu, these micro-clusters undergo a significant, pathogenic expansion. This observation aligns with historical models where small endocrine clusters serve as primary indicators of β-cell neogenesis, a phenomenon previously documented during during pregnancy^26^, insulin-resistance^27^, impired glucose tolerance and new-onset T2D^28^.

This size-dependent expansion is further defined by a profound developmental bias, where nascent EECs are α-cell-predominant. By leveraging the multiplexed capabilities of IMC, we were able to move beyond β-cell counts to resolve the complete endorine cell types of these micro units. We report here the first evidence that human EECs encompass all four major endocrine lineages— α,β,δ, and PP cells—and that each population expands significantly during both prediabetic and T2D stages (Figure 3). However, our quantitative mapping uncovers a critical compositional bias: α-cells serve as the predominant cell type within the extra-islet compartment, with over 65% of EECs existing as “pure” single-hormone units, most of which are glucagon-positive. We excluded ghrelin-producing ϵ-cells from this analysis as they remain extremely scarce in the adult human pancreas^29^. These findings are supported by parallel investigations from our group, which have resolved ductal-associated endocrine remodeling and implicate the ductal epithelium as the most likely source for these nascent, α-cell-rich units (in preparation). This α-cell predominance within the neogenic pipeline may provide a structural mechanism for the systemic glycemic regulation failure observed in T2D. While the pancreas attempts an active regenerative response to metabolic stress, the process is fundamentally skewed toward the glucagon-producing lineage rather than the restoration of functional β-cell mass. This may ultimately contribute to the hyperglucagonemia frequently observed in early diabetes ^14-16,30^, as these clusters expand without the regulatory paracrine “brake” normally provided by β-cell insulin or δ-cell somatostatin^21,22,31^.

### Medium and large islets in T2D progression: density loss and phenotypic remodeling

Medium and large islets (>100 μm) represent the primary structural units of human glycemic stability, yet they undergo catastrophic and disproportionate attrition during the transition to T2D. In this study, these medium and large islets sequester over 90% of total β-cell area, consistent with a previous study which also revealed that relatively larger units accounts for a majority of total β-cell mass ^3^. Our findings of significant density loss in these modules—detectable as early as in prediabetes—extend the seminal observations of Kilimnik et al. regarding large-islet vulnerability in T2D ^32^. Notably, we identified a negative correlation between medium and large islet frequency and donor HbA1c, proving that their preservation is the histopathological prerequisite for normoglycemia.

Beyond depletion in frequency, medium and large islets—the functional “pillars” of glycemic stability—undergo profound cellular compositional and architectural remodeling. In PreD, insulin expression intensity per islet is elevated, aligning with compensatory hyperinsulinemia to counter insulin resistance ^2,33,34^. However, by the onset of overt T2D, the β cell ratio within these islets declines markedly, a shift consistent with dysregulated glycemia. Concomitantly, these islets exhibit progressive structural deterioration during T2D progression, manifested as increased eccentricity, reduced solidity and extent, expanded intra-islet “hollows” (predominantly amyloid deposits), and blurring of boundaries with surrounding exocrine tissue. Because intra-islet paracrine signaling and electrical coupling are fundamental to glycemic regulation ^21,22,31^, this size-selective structural decay physically fragments the endocrine tissue. Such disintegration likely severs critical paracrine coupling circuits, thereby further aggrevates the insulin secretion failure. In particular, the amyloid deposits are harmful to islet cells^35^, which may contribute to islet density loss.

tSNE clustering resolved five discrete populations whose distribution shifted progressively with disease stage: ND islets concentrated in Populations I–III (insulin/β-cell-dominant), while PreD and T2D islets enriched in Populations IV–V (glucagon/α-cell-dominant). The β-to-α cell ratio across this continuum formed an asymmetric bell-shaped trajectory, peaking at Population III (highest β-cell proportion) and declining sharply thereafter. Notably, Population III was most abundant in PreD and in a subset of ND donors with elevated total and LDL-cholesterol, aligning with prior reports of compensatory β-cell expansion during insulin resistance ^36^. This lipid-associated shift in normoglycemic individuals suggests early compensatory remodeling precedes overt hyperglycemia, with PreD sustaining this state before decompensation in T2D (Figure 7).

### Stability of small islets: a divergent pathological fate

A notable finding of this study is the remarkable stability of small islets (30–100 µm in diameter), which evade the size-selective attrition that defines T2D pathology. In contrast to the profound density loss observed in medium and large islets, small islet frequency and their internal β-and α-cell ratios remained statistically unchanged across ND, PreD, and T2D stages. This numerical persistence implies intrinsic resilience to metabolic stressors—such as glucolipotoxicity and amyloid deposition—that drive fragmentation and loss of larger islets.This resilience extends to architectural integrity. Whereas medium and large islets display early morphological deterioration (increased eccentricity and reduced solidity) already in PreD, small islets preserve their compact, circular morphology until the late decompensated phase of T2D. These observations stand in striking contrast to T1D, where small endocrine objects (EOs) are among the earliest targets of immune-mediated destruction^11^. Together, our data on small islets and EECs reveal a fundamental divergence between T1D and T2D: in T2D, smaller endocrine units emerge as the more robust compartment, with EECs even exhibiting expansion and active (albeit α-cell-biased) neogenesis. This size-dependent differential vulnerability highlights distinct pathophysiological mechanisms between autoimmune and metabolic forms of diabetes and suggests that small islets may serve as a relatively preserved reservoir of endocrine function in T2D.

### Limitations

Despite its strengths, IMC provides static data, limiting insights into temporal dynamics; PreD samples partially mitigate this, but longitudinal models (e.g., humanized mice or live human pancreatic slices) could offer further clarity. The antibody panel, while comprehensive, lacks the molecular resolution of spatial transcriptomics, which could uncover underlying mechanisms. The slower image acquisition in IMC restricted our analysis to pancreatic tail sections, though prior studies suggest islet architecture is consistent across pancreas^25^.

## Conclusion

This study provides a comprehensive, high-resolution portrait of human islet heterogeneity and plasticity across the continuum of T2D progression, uncovering a multifaceted failure of glycemic homeostasis driven by size-dependent islet fates and a pervasive α-cell bias (Figure 8). Medium and large islets—the principal reservoirs of β-cell mass—undergo early density loss, architectural disintegration, and phenotypic remodeling, transitioning from initial lipid-driven compensatory expansion to irreversible decompensation marked by β-cell decline and α-cell dominance. In parallel, smaller endocrine units display remarkable resilience: small islets preserve both numerical density and structural integrity, whereas extrainsular endocrine cells (EECs) undergo expansion driven by active, predominantly α-cell-directed neogenesis—a process evident as early as the normoglycemic stage in individuals with elevated lipids. The consistent skew toward glucagon-producing cells—across both neogenesis and remodeling of established islets—suggests a fundamental reprogramming of endocrine fate under metabolic stress. These observations reinforce the longstanding bihormonal hypothesis of diabetes pathogenesis, in which hyperglucagonemia emerges as one central driver of hyperglycemia rather than a mere byproduct of insulin deficiency^16,37,38^. By illuminating the dynamic and heterogeneous responses of human islets to metabolic challenge, our work underscores the need for therapeutic strategies that extend beyond β-cell preservation to address α-cell bias, lipid-mediated stress, and structural integrity of functional islet units. Such approaches may offer new opportunities to halt or reverse the progression of type 2 diabetes.

**Figure 8.**
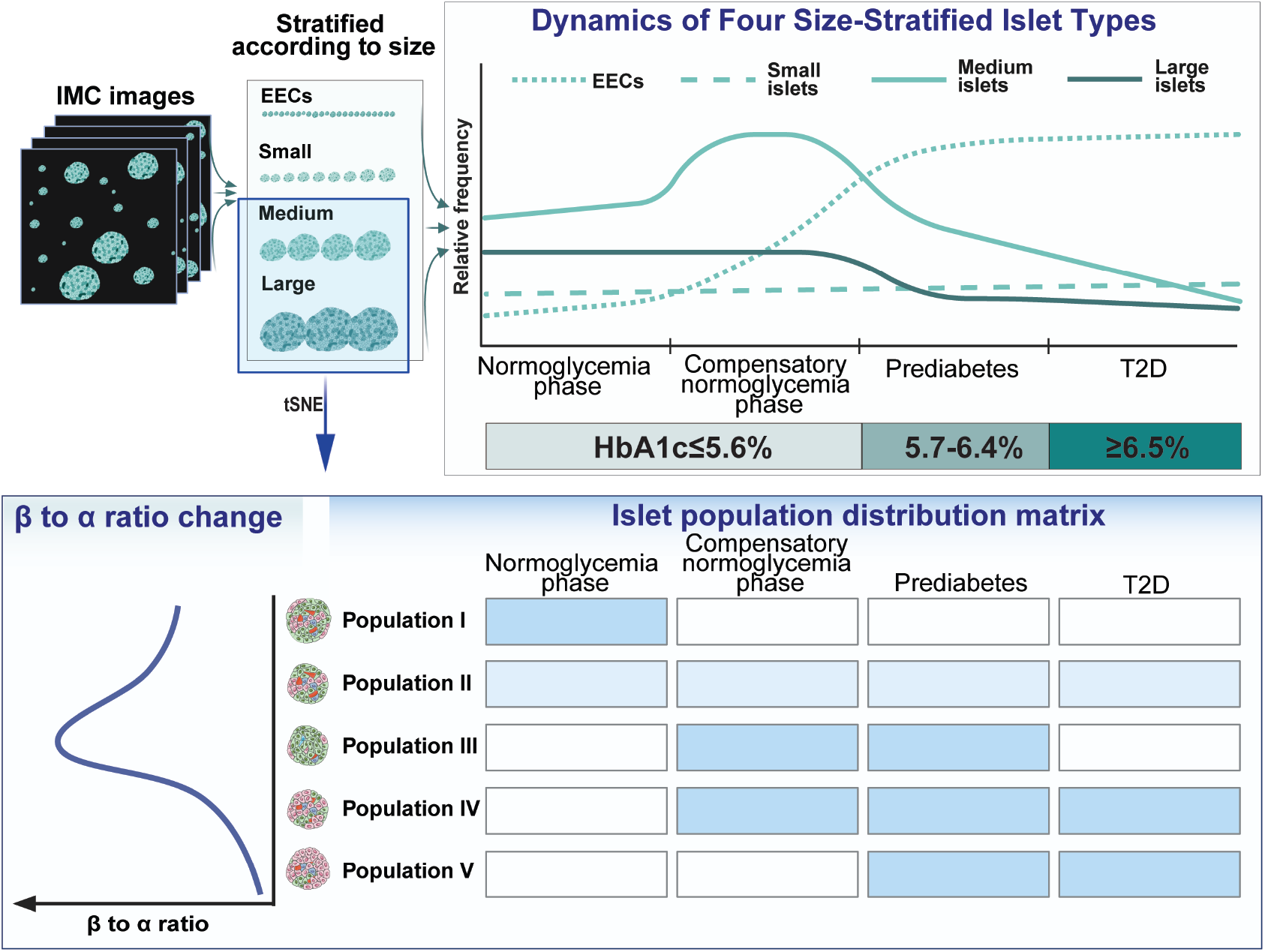
Schematic summary of size-dependent remodeling of the human islet landscape during T2D progression. Abundant EECs are constitutively present in the human pancreas; their expansion initiates as early as the normoglycemic compensatory phase and persists throughout T2D progression. Small islets exhibit relative resilience, with only subtle changes in density during T2D development. In contrast, the density of medium and large islets correlates significantly with glycemic elevation, identifying them as the primary histopathological determinants of systemic glycemic stability. Furthermore, medium and large islets undergo pronounced remodeling of cellular composition: tSNE phenotyping clustered these islets into five distinct populations, revealing a β-cell-dominant compensatory phenotype and an α-cell-dominant diabetic phenotype.

## Acknowledgement

This work was supported by National Key Research and Development Program of China (2024YFA1109002 to S.W.), National Natural Science Foundation of China (82570940 to R.L., 82070805 to S.W.), Tianjin Municipal Human Resources and Social Security Bureau (XB202011 to Human Islet Resource Center, Tianjin), Tianjin Municipal Health Commission (TJWJ2024MS016 to R.L.), Tianjin Key Medical Discipline Construction Project (Grant No. TJYXZDXK-3-013B to L.Z), Chinese Institutes for Medical Research-Beijing to H.R., National Key Research and Development Program of China (2020YFA0803700 to S.W.), and the Tianjin Key Medical Discipline (Specialty) Construction Project (TJYXZDXK-3-011B).

## Methods

### Human organ donors

Thirty paraffin sections of pancreatic tissue samples were selected from our pancreatic tissue bank (Human Islet Resource Center, Tianjin First Central Hospital, China). The pancreases were obtained from organ donors who provided informed consent for research purposes, in accordance with the regulations in China. The study protocol was approved by the Ethical Committee of Tianjin First Central Hospital (Review No.: 2016N073KY).

The samples comprised 10 non-diabetic (ND) donors (HbA1c < 5.7%, no history of diabetes), 8 prediabetic donors (HbA1c between 5.7% and 6.4%), and 12 type 2 diabetic (T2D) donors (HbA1c > 6.4% or a known T2D history) (Table S1). Each group was matched for age and body mass index (BMI) (Table S2). The experimental workflow and data analysis flowchart are presented in Figure 1.

### Antibody conjugation and titration

Validated antibodies (representative IF images were showen in Figure S1) were conjugated to lanthanide metals using the MaxPar X8 Multimetal Labeling Kit (Standard BioTools), following the manufacturer’s instructions. Antibodies were diluted with Antibody Stabilizer containing 0.05% NaN3 (CANDOR® Bioscience, 131050). Serially diluted antibodies were applied to separate sections of the human pancreas to determine the optimal concentrations (Table S3).

### IMC staining

The tail sections of the pancreas were stained using the full antibody panel (Table S4). The slides were incubated in a dry oven at 70 °C for 1 hour, then deparaffinized in fresh xylol and rehydrated through a graded alcohol series. Antigen retrieval was conducted in a microwave oven at 95 °C for 15 min, followed by natural cooling for 20 minutes in Tris-EDTA buffer (pH 9.2). The slides were washed sequentially with ddH2O and DPBS, then permeabilized with 0.5% Triton-100 in DPBS for 15 min and blocked with 3% BSA in DPBS for 45 minutes at room temperature. The slides were incubated overnight at 4 °C with an antibody cocktail (Table S4) containing all the antibodies. The next day, the slides were washed twice in DPBS containing 0.2% Triton-100 and twice in DPBS, then counterstained with the DNA stain Ir for 45 min at room temperature. After washing with ddH2O, the slides were air-dried for 20 min before image acquisition.

### IMC image acquisition

IMC images were acquired at a laser frequency of 200 Hz, following daily tuning and the manufacturer’s protocols from Standard BioTools. The regions of interest (ROIs) were selected based on the locations of islets identified in the H&E staining. Each ROI measured 800 µm × 800 µm and aimed to include as many islets as possible, with at least one islet per area (Figure S2). On average, 9 ROIs (range: 5–12) were selected per slide. The MCD files were converted to TIFF images using Standard BioTools’ MCD viewer and processed for islet and cell segmentation to extract the data. Representative IMC images were shown in Figure S3.

### Islet segmentation

Binary masks for insulin (INS), glucagon (GLU), somatostatin (SST), and pancreatic polypeptide (PP) hormone were summed to generate a composite islet mask. Morphological holes within the composite mask were filled using the *imfill* function using MATLAB (MathWorks, Natick, MA, USA). Connected component analysis was performed using the *regionprops* function with 8-connected neighborhood connectivity. Objects with cross-sectional areas smaller than xx um2 (parameter determined based on expected minimum islet size) were excluded.

### Cell segmentation

CellProfiler 4.2.1 (Kamentsky et al., 2011) was used to define cell masks. We employed DNA-Iridium (Iridium 191 and 193) as nuclear markers to define primary objects. Primary objects with an area of less than 5 pixels and more than 15 were filtered out. CD99, INS, GCG, SST, and PP staining were thresholded and then combined to serve as the membrane marker for cell image. We employed Distance-N method to identify the secondary objects based on Cell Image and primary objects. Specifically, primary objects recognized in the previous step were expanded with guidance of the cell membrane probability map, with maximum expansion constrained as 5 pixels. Two independent cell masks for each image set were generated by two different investigators. Lastly, we used Cell mask in MATLAB software (MathWorks, Natick, MA, USA) to measure mean protein expression intensities for each channel and each cell in the original IMC images post pre-processing steps. The x and y coordinate of all cells in the image were also recorded.

### Cell type classification

Cell types were defined based on the occupancy ratio of binary masks, with a 20% threshold applied to distinguish positive staining. Specifically, cells with ≥20% occupancy in more than one binary mask (INS, GLU, SST, PP) were classified by their highest relative occupancy ratio among the four hormones. This approach minimized repetitive counting and ensured unambiguous cell-type designation in islet cell type composition analyses.

### Islet features

For each islet, Cell number is defined as total cell numbers in the islet. Area is defined as total pixel counts (each pixel is 1 µm^2^). Cell number is defined as total cell numbers in the islet. Diameter is defined as the length in pixels of the major axis of the ellipse. Solidity is defined as the proportion of the pixels in the convex hull that are also in the region. Extent is defined as the ratio of pixels in the region to pixels in the total bounding box. Eccentricity is the ratio of the distance between the foci of the ellipse and its major axis length. The value is between 0 and 1. (0 and 1 are degenerate cases. An ellipse whose eccentricity is 0 is actually a circle, while an ellipse whose eccentricity is 1 is a line segment.) Area, diameter, solidity, extent and eccentricity are defined as the *Area, MajorAxisLength, Solidity, Extent* and *Eccentricity* from regionprops function. Type is defined based on the type of patient, ND PreD or T2D. Quantitative Analysis of Protein composition metrics were calculated to characterize intra- and extra-islet expression patterns: Intra-islet Mean Protein Intensity (Mean_protein_dist_1): The mean fluorescence intensity of the target protein within the boundaries of identified islets. Extra-islet Mean Protein Intensity (Mean_protein_dist_2): The mean fluorescence intensity of the target protein in the surrounding tissue excluding islet regions. Protein includes INS, NGN3, CD34, SOX9, ARX, GLU, PP, PDX1, SST, SYN, CK19, P_H3, UCN2, SMA, KI67, NKX6.1, ALDH, GHRELIN, CD99 and BACT. Intra-islet Protein Mask Ratio (Mean_protein_mask_dist_1): The proportion of the islet area occupied by the protein’s binary mask. Extra-islet Protein Mask Ratio (Mean_protein_mask_dist_2): The proportion of non-islet tissue occupied by the protein’s binary mask. Protein mask includes ARX, CD34, CD99, CK19, GLU, INS, KI67 NKX6.1, PP, SST and SMA. Analyses were performed using custom MATLAB scripts.

### Statistics

For group comparisons, we systematically used one way ANOVA accompanied by Fisher’s LSD test for multiple comparisons. For correlation analysis, Pearson’s correlation coefficient method was used.

Overall, the p value significance threshold was defined as 0.05. In text and figures, n represents the number of donors, ROIs, islets, or cells (as indicated in figure legends). In histograms, bars represent the means and error bars represent standard errors of the mean.

### Data and software ability

The accession number for the IMC data reported in this paper (including image stacks, single-cell data and islet-level data) can be obtained in Mendeley Data when the manuscript is published. Custom code (Matlab) generated by the Huixia Ren lab for IMC data preprocessing and image segmentation will be available online when the manuscript is published; MCD viewer for image reconstruction can be downloaded at https://www.standardbio.com/products/software. Cell profiler for cell segmentation is available at https://cellprofiler.org/.

