## Supplementary figures for "A spatial-temporal atlas of human islet pathophysiology identifies a size-dependent trajectory from compensation to decompensation"

**Figure S1. Representative images of each antibody in testing immunofluorescence staining of pancreas tissue.**

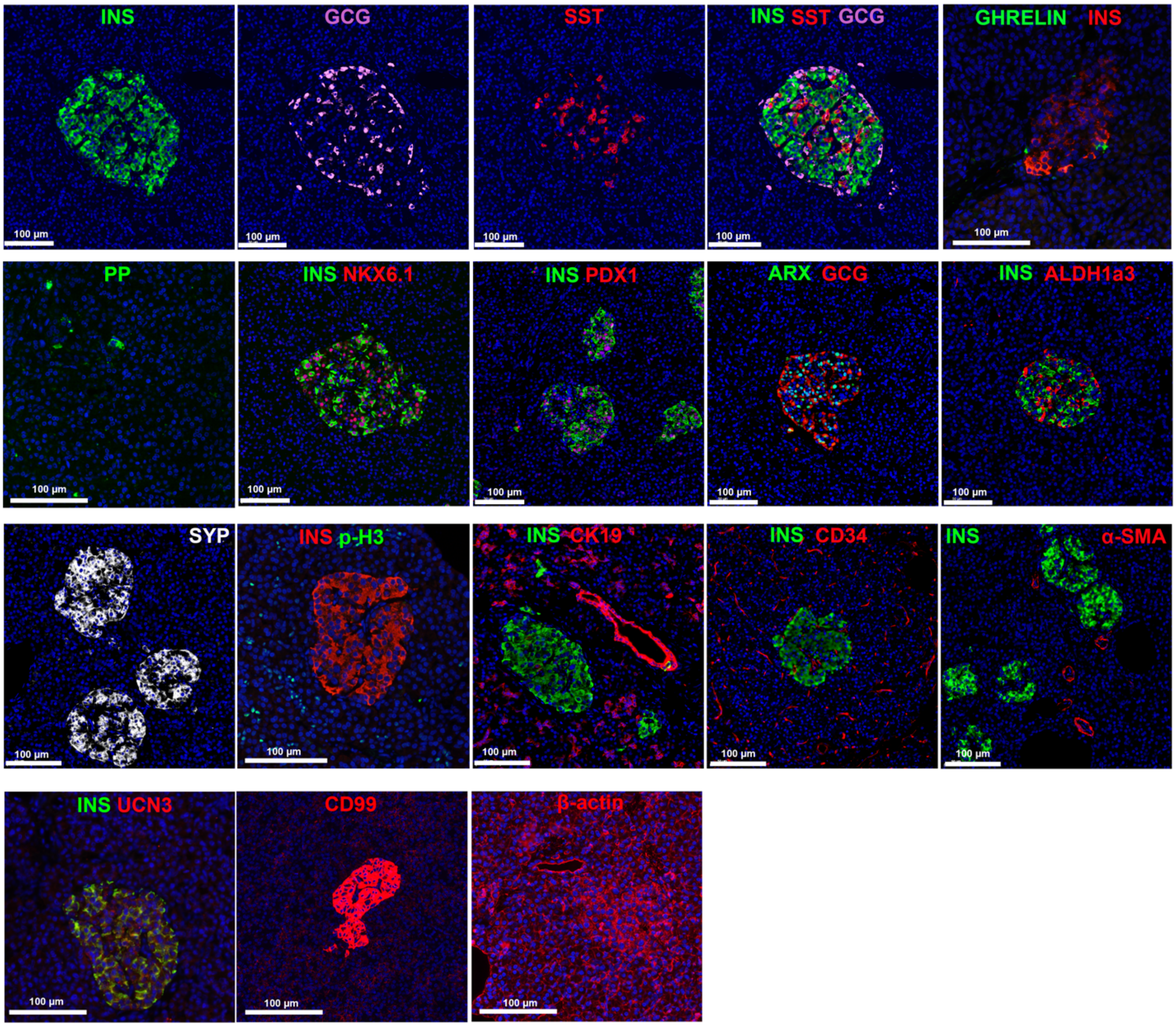

**Figure S2. Selection of regions of interest (ROIs).** H&E was initially on each slide, followed by IMC staining on sequential sections. ROIs were chosen based on the location of islets identified in the H&E or immunofluorescence staining results. Each ROI is 800 µm ×800 µm and aimed to include as many islets as possible, with a minimum of one islet per ROI.

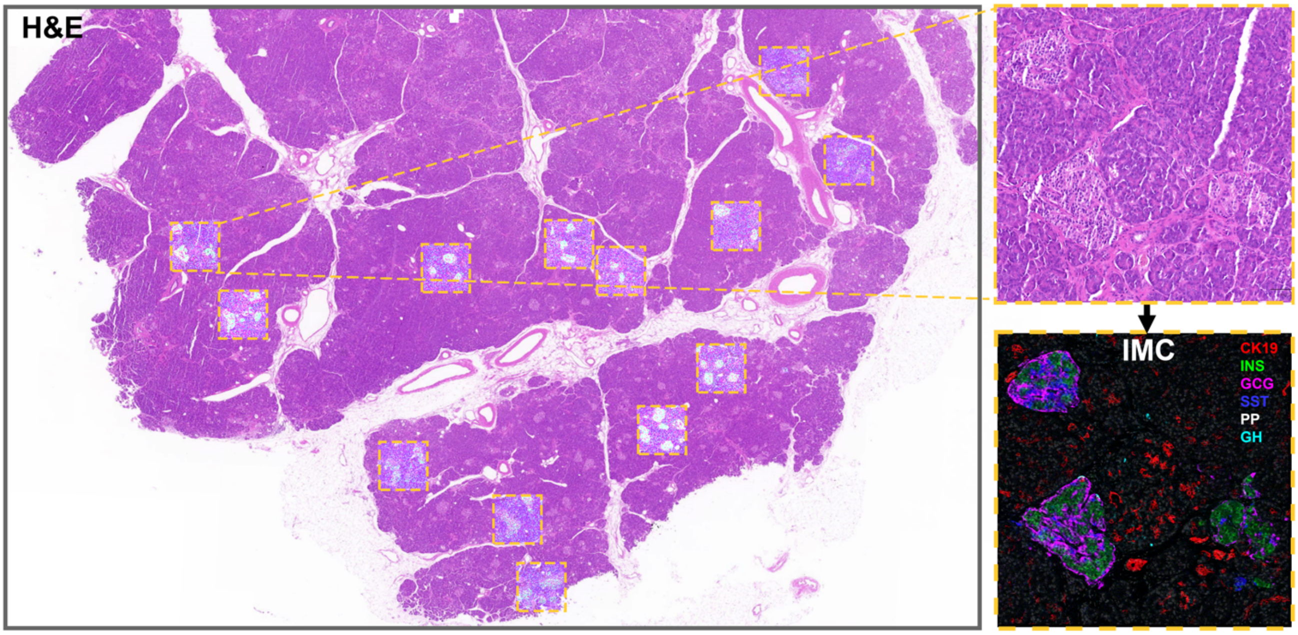

**Figure S3. Representative images of each isotope-labeled antibody in the IMC staining of pancreas tissue. (Green: marker protein; Blue: DNA)**

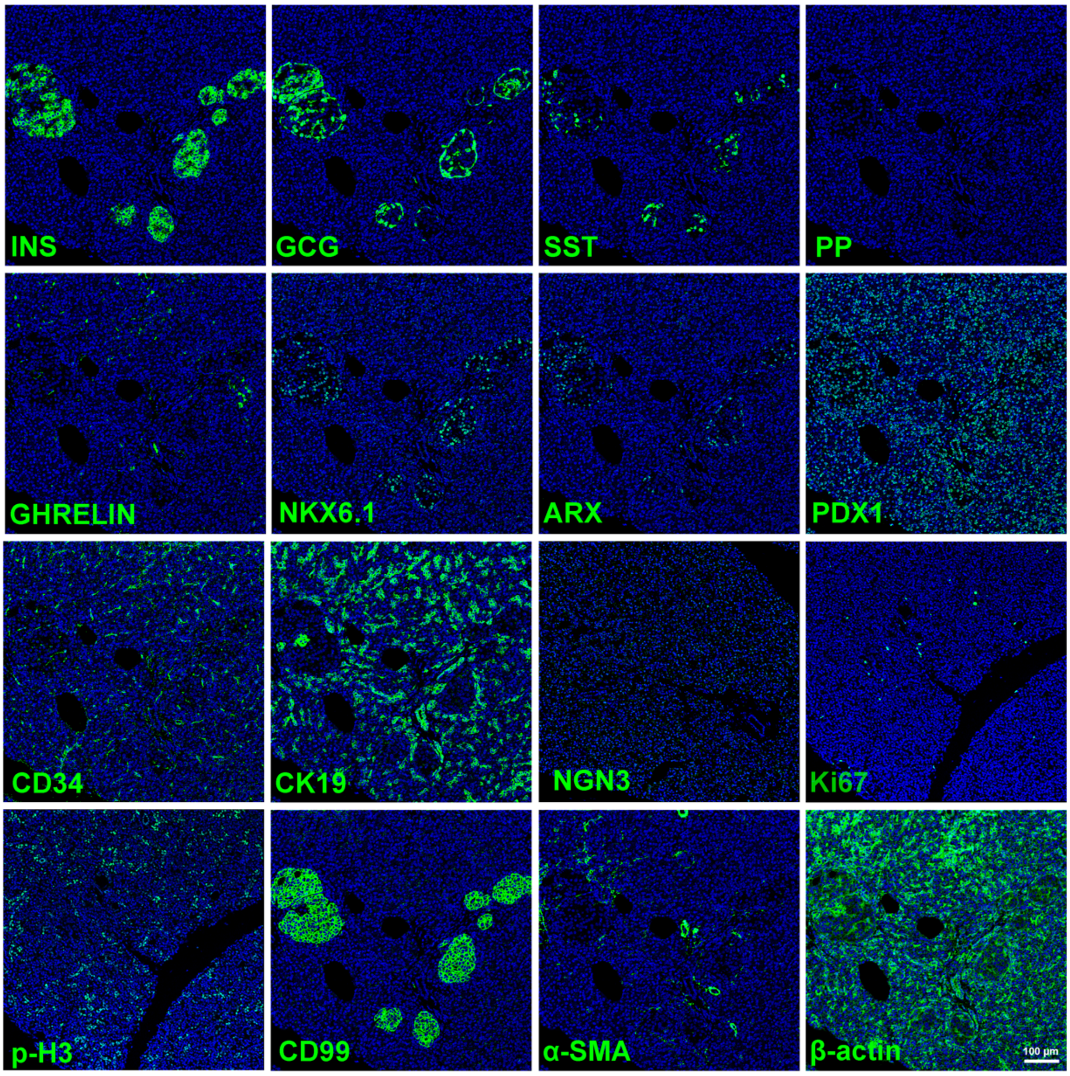

**Figure S4. Projection of islet area to diameter. The value labeled on the top of the violin column is the mean of the diameters of islets in each area bin.**

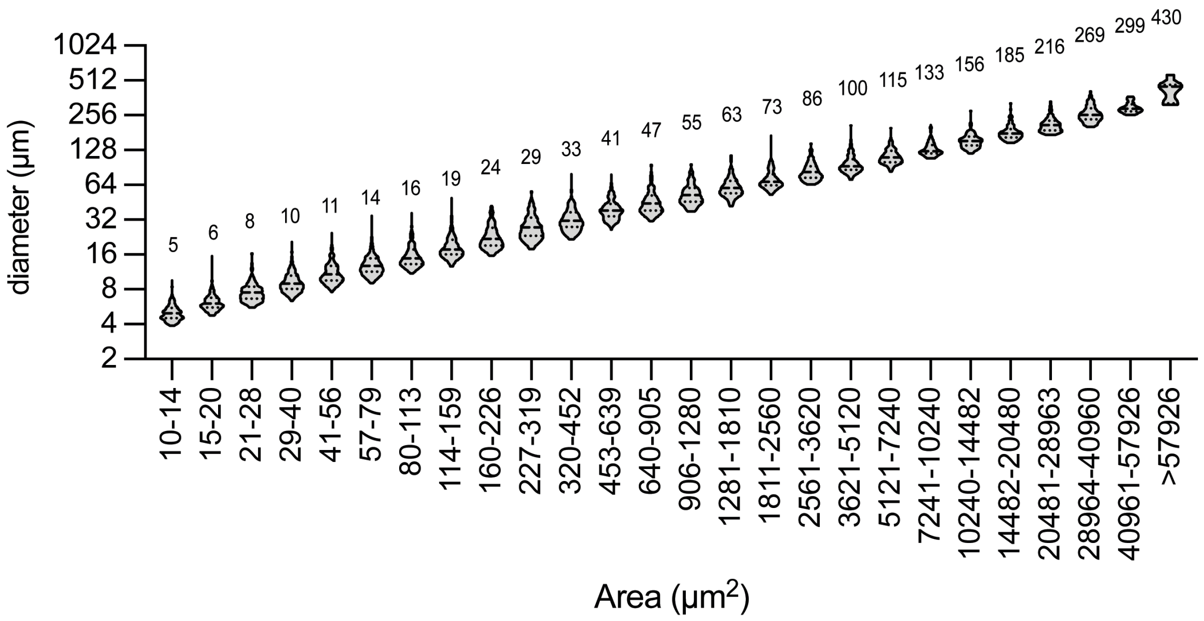


**Figure S5. Percentage of EECs with only one hormone expression.**

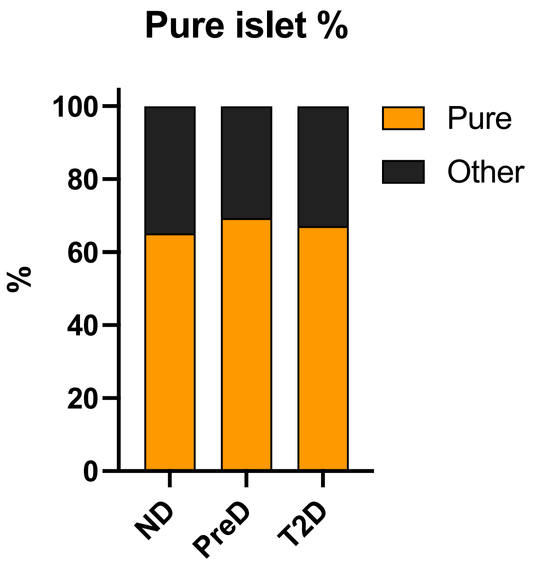

**Figure S6. Representative IMC images of islet hole (intra-islet amyloid deposits) in T2D subjects.** Top layer: H&E staining images; bottom layer: IMC images. Green: insulin; Red: glucagon; Blue: DNA; Scale bar: 100 µm.

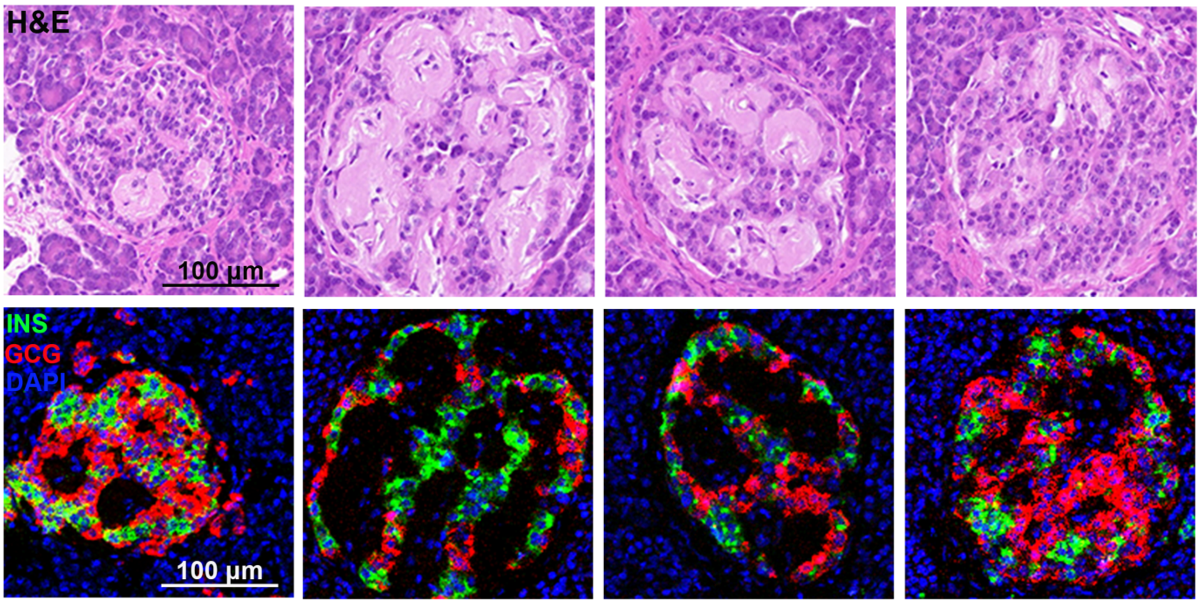
