## Supplementary tables for "A spatial-temporal atlas of human islet pathophysiology identifies a size-dependent trajectory from compensation to decompensation"

| Donor No. | Type | Age (years) | Gender | BMI (kg/m2) | HbA1c (%) | C-Peptide (ng/ml) | Diabetes duration (years) | TG (mM) | CH (mM) | LDL-C (mM) |
|---|---|---|---|---|---|---|---|---|---|---|
| **HP025** | **ND** | 37 | male | 21.72 | 5.5 | 3.85 | / | 0.53 | 3.14 | 1.67 |
| **HP028** | **ND** | 44 | male | 22.49 | 4.3 | / | / | 0.39 | 1.36 | 0.68 |
| **HP035** | **ND** | 45 | male | 27.78 | 5.6 | 12.77 | / | 0.9 | 2.51 | 1.4 |
| **HP039** | **ND** | 49 | female | 30.82 | 5.5 | 2.25 | / | 0.46 | 4.97 | 2.32 |
| **HP045** | **ND** | 58 | male | 26.12 | 5.3 | 2.6 | / | 2.17 | 3.79 | 1.8 |
| **HP046** | **ND** | 52 | male | 20.76 | 5.1 | 3.37 | / | / | / | / |
| **HP048** | **ND** | 57 | male | 22.49 | 5.5 | 5.14 | / | 0.3 | 2.89 | 0.86 |
| **HP051** | **ND** | 30 | male | 23.66 | 5.5 | 4.93 | / | 1.23 | 3.41 | 1.22 |
| **HP052** | **ND** | 53 | female | 20.96 | 5.5 | 1.94 | / | / | / | / |
| **HP111** | **ND** | 47 | male | 27.34 | 5.4 | 14.8 | / | 2.84 | 4.05 | 2.28 |
| **HP034** | **PreD** | 43 | female | 21.48 | 6.1 | 3.71 | / | 0.62 | 1.48 | 0.71 |
| **HP050** | **PreD** | 60 | male | 22.86 | 5.9 | 2.86 | / | 1.13 | 3.05 | 1.54 |
| **HP061** | **PreD** | 53 | male | 23.51 | 6 | 29.96 | / | / | / | / |
| **HP094** | **PreD** | 26 | male | 19.59 | 6.4 | 2.86 | / | 1.09 | 1.93 | 0.81 |
| **HP110** | **PreD** | 65 | female | 21.48 | 5.8 | 3.87 | / | 0.93 | 2.82 | 1.18 |
| **HP112** | **PreD** | 44 | male | 23.66 | 5.9 | 5.91 | / | 0.91 | 4.49 | 2.96 |
| **HP113** | **PreD** | 44 | male | 29.39 | 6.1 | 6.46 | / | 1.59 | 4.95 | 3.72 |
| **HP127** | **PreD** | 20 | male | 20.98 | 5.7 | 2.59 | / | 1.42 | 4.18 | 2.52 |
| **HP012** | **T2D** | 63 | male | 25.95 | 7.2 | 0.311 | 10 | 0.21 | 1.02 | 0.44 |
| **HP014** | **T2D** | 49 | male | 29.40 | 6.9 | 6.22 | 10 | 1.14 | 1.97 | 1.09 |
| **HP017** | **T2D** | 49 | male | 24.20 | 7.6 | / | / | / | / | / |
| **HP018** | **T2D** | 47 | male | 32.40 | 6.7 | 5.13 | / | 1.07 | 4.04 | 2.36 |
| **HP019** | **T2D** | 44 | male | 24.20 | 6.1 | 9.41 | 0.08 | 1.25 | 4.32 | 2.5 |
| **HP027** | **T2D** | 59 | male | 22.49 | 9.6 | 4.87 | / | 0.61 | 4.88 | 2.54 |
| **HP037** | **T2D** | 42 | male | 30.86 | 6.3 | / | 5 | 1.87 | 3.36 | 1.67 |
| **HP038** | **T2D** | 55 | male | 22.59 | 6.8 | 5.47 | / | 1.71 | 3.08 | 1.66 |

| | | | | | | | | | | |
|---|---|---|---|---|---|---|---|---|---|---|
| **HP053** | **T2D** | 45 | male | 26.23 | 6.5 | 5.63 | / | / | / | / |
| **HP054** | **T2D** | 65 | male | 23.93 | 7.5 | / | 5 | 0.85 | 2.75 | 1.32 |
| **HP055** | **T2D** | 64 | female | 23.43 | 8.9 | 1.71 | / | 1.24 | 3.49 | 2.04 |
| **HP105** | **T2D** | 57 | female | 24.97 | 5.8 | 7.71 | 30 | 1.37 | 3.55 | 2.18 |

TG: Triglycerides
CHO: cholesterol
LDL-C: low-density lipoprotein cholesterol

**Table S2. Donor information in ND, PreD, T2D groups.**

| Groups | N | F/M | Age (year) | BMI (Kg/m$^2$) | HbA1c (%) | C-Peptide (ng/ml) | TG (mM) | CH (mM) | HDL-C (mM) | LDL-C (mM) |
|---|---|---|---|---|---|---|---|---|---|---|
| ND | 10 | 2/8 | 47.20 ± 8.74 | 24.41 ± 3.41 | 5.32 ± 0.39 | 5.74 ± 4.72 | 1.10 ± 0.93 | 3.27 ± 1.08 | 1.00 ± 0.45 | 1.53 ± 0.61 |
| PreD | 8 | 2/6 | 44.38 ± 15.48 | 22.87 ± 2.97 | 6.00 ± 0.22 | 7.28 ± 9.28 | 1.10 ± 0.33 | 3.27 ± 1.32 | 0.99 ± 0.39 | 1.92 ± 1.16 |
| T2D | 12 | 2/10 | 53.25 ± 8.28 | 25.89 ± 3.28 | 7.16 ± 1.12$^{a,b}$ | 5.16 ± 2.77 | 1.13 ± 0.49 | 3.25 ± 1.13 | 0.78 ± 0.37 | 1.78 ± 0.68 |

Data are shown as mean ± SD. A: $^{a}P < 0.001$, T2D vs ND; $^{b}P < 0.01$, T2D vs PreD

**Table S3. Antibody information.**

| Antibodies | Source | Cat. | Expression | Metal label | Dilution |
|---|---|---|---|---|---|
| **Insulin** | Abcam | ab63820 | β | 141Pr | 1:400 |
| **NKX6.1** | Novus | NBP1-49672 | β | 169Tm | 1:200 |
| **PDX1** | CST | D59H3 | β | 154Sm | 1:25 |
| **UCN3** | Sigma | HPA038281 | β | 165Ho | 1:200 |
| **Glucagon** | Abcam | ab10988 | α | 151Eu | 1:400 |
| **ARX** | R&D | AF7068 | α | 149Sm | 1:200 |
| **Somatostatin** | Santa Cruz | sc-55565 | δ | 159Tb | 1:1600 |
| **Pancreatic Polypeptide** | Abcam | ab77192 | PP | 153Eu | 1:200 |
| **Synaptophysin** | Abcam | ab32127 | Endocrine | 160Gd | 1:50 |
| **ALDH1a3** | Abcam | NBP2-15339 | Progenitor/Dedifferentiated | 170Er | 1:800 |
| **NGN3** | R&D | AF3444 | Progenitor/Dedifferentiated | 142Nd | 1:100 |
| **SOX9** | Abnova | H00006662-M02 | Progenitor/Dedifferentiated | 147Sm | 1:50 |
| **CK19（cytokeratin 19 antibody）** | Novus | NB100-687SS | ductal | 161Dy | 1:200 |
| **CD34** | Abcam | ab81289 | Endothelial | 146Nd | 1:50 |
| **α-SMA** | Sigma | A5228 | Stromal/Stellate | 167Er | 1:800 |
| **Ki67** | Abcam | ab15580 | Proliferating | 168Er | 1:25 |
| **Histone H3 Phospho (Ser28)** | BioLegend | 641002 | Proliferating | 163Dy | 1:400 |
| **GHRELIN** | Santa Cruz | sc-293422 | Epsilon | 173Yb | 1:100 |
| **CD99** | R&D | AF3968 | Cell Member marker | 175Lu | 1:100 |
| **β-actin** | CST | 3700s | Cell Member marker | 176Yb | 1:200 |
